# VAMP2 AND SYNAPTOTAGMINS ARE RELATIVELY IMMOBILE ON CHROMAFFIN GRANULE MEMBRANES: IMPLICATIONS FOR MEMBRANE FUSION AND FUSION PORE EXPANSION

**DOI:** 10.1101/2021.02.20.432015

**Authors:** Prabhodh S. Abbineni, Joseph S. Briguglio, Edwin R. Chapman, Ronald W. Holz, Daniel Axelrod

## Abstract

Although many of the proteins of secretory granules have been identified, little is known about their molecular organization and diffusion characteristics. Granule-plasma membrane fusion can only occur when proteins that enable fusion are present at the granule-plasma membrane contact. Thus, the mobility of granule membrane proteins may be an important determinant of fusion pore formation and expansion. To address this issue, we measured the mobility of (fluorophore-tagged) vesicle associated membrane protein 2 (VAMP2), synaptotagmin 1 (Syt1), and synaptotagmin 7 (Syt7) in chromaffin granule membranes in living chromaffin cells. We used a method that is not limited by standard optical resolution. A bright flash of strongly decaying evanescent field (∼80 nm exponential decay constant) produced by total internal reflection (TIR) was used to photobleach GFP-labeled proteins in the granule membrane. Fluorescence recovery occurs as unbleached protein in the granule membrane distal from the glass interface diffuses into the more bleached proximal regions, thereby enabling the measurement of diffusion coefficients. The studies revealed that VAMP2, Syt1, and Syt7 are relatively immobile in chromaffin granules membranes with diffusion constants of ≤ 3 × 10^−10^ cm^2^/s. Utilizing these diffusion parameters and the known density of VAMP2 and Syt 1 on synaptic vesicles, we estimated the time required for these proteins to arrive at a nascent fusion site to be tens of milliseconds. We propose that the mobilities of secretory granule SNARE and Syt proteins, heretofore unappreciated factors, influence the kinetics of exocytosis and protein discharge.

**Significance Statement:** In eukaryotic cells, secretory vesicles fuse with the plasma membrane to secrete chemical transmitters, hormones and proteins that enable diverse physiological functions including neurotransmission. Fusion proteins need to be assembled at the fusion site in sufficient number in order to enable membrane fusion. However, the diffusion characteristics of fusogenic proteins on secretory vesicles remained unknown. Here we used a novel method not limited by standard optical resolution to measure the diffusion of VAMP2 and synaptotagmins on chromaffin granule membranes. We found they have limited mobility. The time required for these proteins to reach the granule-plasma membrane contact site suggests that their limited mobility likely influences the kinetics of membrane fusion and subsequent fusion pore expansion.

## Introduction

Enormous progress has been made in understanding the protein interactions and events leading to fusion of secretory granules and synaptic vesicles with the plasma membrane. Nevertheless, the final events leading to fusion remain uncertain. Granules must move into intimate contact with the plasma membrane in order for granule membrane and plasma membrane proteins to interact to initiate the fusion reaction. These proteins include the granule SNARE protein VAMP2 (synaptobrevin-2) and the Ca^2+^ sensor synaptotagmin and the plasma membrane acceptor complex composed of the t-SNARES, SNAP25 and syntaxin (1, 2). It has not been established when these interactions occur. On the one hand, the initial interactions may create a stable intermediate that awaits a Ca^2+^ signal to pull the membranes together to initiate fusion (3). Alternatively, the constantly jittering granules (4, 5) only interact with the plasma membrane milliseconds or less before fusion. Indeed, it has been suggested that the strongly energy-favorable interactions of opposing SNARE proteins may not allow a stable intermediate (2); once engaged, the interactions proceed rapidly to fusion.

Close approximation of the granule and plasma membrane does not guarantee productive protein interactions. A kinetic factor in fusion that has not considered is the speed with which proteins access the ultimate fusion site. This is especially true in chromaffin cells in which exocytosis is widely distributed over the entire plasma membrane (6, 7) and does not occur at readily identifiable fusion sites (8). These considerations raise the possibility that protein mobility in the two opposing membranes could play a role in determining the rate of granule fusion. For example, if the proteins are highly mobile in both membranes, a close approach of the granule and plasma membranes would have a significant probability of enabling interaction of the cognate proteins. Alternatively, if the proteins are immobile, nanometer distance apposition of the two membrane would only result in fusion if by chance the cognate proteins are already present and properly oriented to enable interaction. In this second scenario, motion of the entire granule may aid interaction of fusogenic granule and plasma membrane proteins and lipids.

In this study we have investigated the mobility of VAMP-2, synaptotagmin1 (syt1) and synaptotagmin7 (syt7) in the secretory (chromaffin) granule membrane in living chromaffin cells. We adapted a method that is not limited by standard optical resolution that we had previously developed to measure the mobility of granule lumenal proteins (9). Granules are bleached in a strongly decaying evanescent field (∼80 nm exponential decay constant) produced by total internal reflection (TIR). Since the diameter of chromaffin granules is ∼300 nm, the high intensity excitation selectively bleaches fluorophore-label protein proximal to the glass interface in the membrane of individual chromaffin granules. Fluorescence recovery can occur as unbleached protein from the more distal surface of the granule diffuses into the proximal bleached regions. This experimental approach, a variation of TIR-fluorescence recovery after photobleaching (TIR-FRAP) (10) introduced for open areas in 1981, is accompanied by a new theoretical and quantitative analysis that takes into account the limited number of total fluorophore molecules on the granule membrane, diffusion in the spherical membrane, granule diameter, the evanescent field depth and the duration of the bleach. In the course of the experiments, we discovered additional experimental factors that had to be considered: light scattering that accompanies TIRF excitation, and reversible bleaching of the GFP-based fluorophores and its relationship to irreversible bleaching. Finally, we consider the influence of VAMP2 and Syt diffusion on initiating fusion and subsequent fusion pore expansion.

## THEORY

In these experiments, a secretory granule (modeled as a 300 nm diameter hollow sphere) is embedded in an evanescent field (created by TIR) with an exponentially-decaying characteristic depth (80 nm) on the same order of size as the sphere’s radius R. There are two versions of the experiment: one for fluorophore-labeled membrane proteins confined to 2D diffusion on the granule surface, and the other for fluorophore-labeled luminal proteins confined to 3D diffusion within the granule interior. The qualitative interpretations and relevant complications for the two versions of experiments are the same; the only difference lies in the quantitative theoretical interpretation.

The experimental protocol for the two versions is the same. A dim “probe” intensity of the evanescent field excites fluorescence, predominately but not exclusively, near the “bottom” of the sphere where the sphere is proximal to the TIR substrate (the coverslip) and the evanescent field is strongest. The total TIR-excited fluorescence from the whole sphere is measured and normalized to unity. Then, the illumination intensity is increased in a single step to a factor of ∼100 higher for a duration of 46 ms (for membrane proteins) or 169 ms (for luminal proteins), which leads to significant bleaching of the fluorophore, especially but not exclusively at the granule bottom. Then after the bleach, the evanescent intensity is reduced to its pre-bleach level, the emitted total fluorescence intensity is tracked vs. time, as unbleached fluorophores diffuse around (or in) the sphere toward a uniform distribution.

In principle, the form of the recovery could be calculated (perhaps numerically) from an explicit differential equation. However, there are several significant real-world factors which make the solution, and the ultimate determination of the diffusion coefficient, somewhat more complicated:

a. The excitation itself is not a pure TIR-produced exponential decay, but contains some fraction of its intensity arising from scattering, originating either from the microscope optics (11, 12) or from refractive index irregularities in the sample itself (13). That scattering fraction can lead to its own bleaching during all phases of the experiment. The actual excitation intensity thereby is a sum of the exponentially-decaying evanescent part plus a z-independent scattering part, estimated here to be an additional ∼20% of the z=0 evanescent intensity.
b. The time scale of the experiments is short, on the order of tens to hundreds of milliseconds. On that time scale, bleaching is not entirely irreversible; some bleached fluorophores return spontaneously to a ground state (e.g., a return from a long-lived triplet state), capable of re-excitation to a fluorescence-producing excited state. This reversible recovery must be distinguished from the diffusion-based recovery.
c. The mechanism of reversible bleaching is not well-known. Possible modes include the following. (1) Reversible and irreversible bleaching are parallel processes, either occurring to an unbleached fluorophore with some fixed likelihood. (2) A reversibly bleached fluorophore is protected against subsequent irreversible bleaching until it returns to the unbleached ground state. (3) Irreversible bleaching can occur only to a fluorophore which is already in a reversibly bleached state. Each of these possibilities would lead to a different dependence of ratio of reversible:irreversible bleach depth as the incident illumination intensity is changed. The question of which mode is correct can be investigated by examining reversible vs irreversible bleach depth under EPI-illumination. (In EPI, the excitation intensity through the entire depth of the granule is constant, and diffusion-related effects would thereby become irrelevant.) Our observations are most consistent with alternative (3): that irreversible bleaching occurs mainly to fluorophores while they are in a reversible bleached state. If experiments had been at a much longer time scale, reversible bleaching would not be evident, but that would be too long to see the diffusion effects that motivate the experiments.
d. During the prebleach and postbleach dim “probe” phase, significant bleaching (either reversible or irreversible or both) can occur, which tends to cause a loss of fluorescence. Because the illumination is z-dependent, even probe phase bleaching occurs non-uniformly around the sphere’s surface or in its interior.
e. During the finite-time duration of the bright “bleach” phase, significant diffusion of the fluorophore can occur, making the t=0 distribution of fluorophores immediately at the end of the bleach phase somewhat poorly-defined.
f. Each granule may be located at a slightly different distance from the TIR/coverslip surface and thereby sees a different evanescent illumination intensity. This variability will manifest as a different t=0 postbleach fluorescence. That variability will affect the postbleach distribution of unbleached fluorophores and ultimately the exact form and time scale of the fluorescent recovery. For this reason, results are grouped and averaged into several separate ranges of bleach depth.

To account for all these complications, we use a custom Monte Carlo-type program (written in IDL and named “diffusionshellsim”) that repeatedly simulates the diffusion as a virtual random walk of a single fluorophore molecule. A simulated membrane protein diffusing on a spherical surface, or within the sphere, is first positioned at a random location in its realm. During its walk, the fluorescence vs time it produces is calculated and recorded from its z-position (the distance from the TIR/coverslip surface), given a z-dependent excitation illumination intensity that includes both evanescent decay and scattering. That molecule is tracked over its entire random walk course vs. time until it irreversibly bleaches. Then another molecule is launched from a new random location. Altogether, a complete simulation consists of summing the fluorescence vs time of tens of thousands of such single molecule tracks.

Here is more detail on this procedure, and how the complications are handled. Time is divided into a large (consistent with an acceptable total computation time) but finite number of time increments (each with a duration of 1 millisecond), out to a simulated time of some t=t_max_. For surface-confined membrane proteins, in each time increment in sequence, two new normally-distributed random numbers (symmetrical around zero) are generated, one to compute the random length (generally <<R) of a single step to a new location and the other its random direction angle from its starting point on the surface. After appropriate geometrical considerations involving rotation matricies, the z position of the new location is calculated. The average of the length-squared of the diffusive step vectors is scaled to be proportional to an input parameter diffusion coefficient D. For volume-confined lumenal proteins, in each time increment in sequence, three new normally-distributed random numbers (symmetrical around zero) are generated, one for each of the three orthogonal dimensions, with the average step size corresponding to a user-specified diffusion coefficient. If the next step places the molecule outside the sphere by some distance *l* from the surface, it is “reflected” back into the sphere to a location along the same radial line and the same distance *l* from the surface.

All of the bleaching effects (b)-(f) above can be incorporated into each incremental step by using additional input parameters relevant to bleaching efficiency. These parameters, and how they are determined, are explained as follows:

1. *The characteristic rate (in sec*^*-1*^*) for reversible bleach recovery*. This rate parameter can be determined directly from experimental data taken on granules with EPI illumination (i.e, subcritical angle) instead of TIR illumination optics using the same bleach-and-probe intensity variation protocol. Since EPI illumination is not z-dependent at the sample, the consequent fluorescence recovery, if any, is entirely due to reversible recovery.
2. *A* “*total bleaching” parameter, proportional to local excitation intensity, that determines the immediately postbleach (t=0) fluorescence*. This parameter can be varied in the simulation program to produce simulated fluorescence at t=0 that matches that of the average of group of similar-bleach-depth experimental runs.
3. *The ratio of probe to bleach illumination, which is known by direct pre-measurements of laser intensity entering the microscope*.
4. *The ratio of the reversible to irreversible bleaching probabilities*. This parameter strongly affects the ratio of the fluorescence at t=0 to the long-time “plateau” fluorescence (measured in practice at t=t_max_) in the EPI illumination experiments (which are insensitive to diffusion). This ratio can be adjusted in the simulation, essentially by trial-and-error, to produce simulated EPI results that correspond the experimentally observed EPI fluorescence ratio for t=0 to t=t_max_.

These input parameters can be expressed as single step probabilities that a molecule will be either left unbleached, reversibly bleached, recovered from prior reversible bleaching, or irreversibly bleached. [Any probability (say, *p*) is realized by uniformly randomly generating a number between 0 and 1 and proceeding appropriately if the number turns out to be less than *p*.] In our preferred mode, irreversible bleaching is possible only if the molecule has already been reversibly bleached in a previous step. If the molecule does not get irreversibly bleached, it “survives” to make the next diffusive step on the sphere, and the process of determining its fluorescence and its bleach status is repeated. If the molecule is irreversibly bleached at any step, its fluorescence vs time history up to that time is added to that of previous molecule fluorescence histories, and then a new molecule is started at a random position. After a preset number of molecules (usually in the tens of thousands) is reached, the final accumulated fluorescence vs time curve is the simulation program’s output.

The overall goal is to determine the one remaining parameter, the diffusion coefficient D. Simulations with the correct bleaching parameters are run for a range of possible D values. The simulated results are then compared to the corresponding experimental TIR data (the average of runs with similar bleaching depths) by calculating the average unweighted chi-square difference between the experimental and simulated curve. The D for which the chi-square value is at a minimum is deemed to be the correct diffusion coefficient.

## RESULTS

### Simulation of diffusion on the surface of a sphere

We utilized two methods that rely on photobleaching with total internal reflection (TIR) excitation to measure the diffusion of fluorophore-tagged VAMP2, Syt1, and Syt7 on chromaffin granule membranes. Following photobleaching with TIR excitation, a gradient of fluorescence is imprinted on the granule membrane (**Fig. 1 A**.), as the evanescent field decay constant (∼ 80 nm) is shorter than the diameter of chromaffin granules (∼ 300 nm). Redistribution of fluorophores into the bleached region by diffusion leads to at least partial dissipation of the gradient (**Fig.1A, compare Immobile and Mobile fluorophores)**. The faster the diffusion, the more rapidly the gradient suffers dissipation. The first method (“bleach depth”) measures the presence of the gradient by measuring the fluorescence intensity of the granules before and after a TIR bleach using two kinds of probe illumination in rapid sequence 1) a low intensity TIR evanescent field that excites the region that will be subsequently bleached or just has been bleached; and 2) an epifluorescence (EPI) illumination probe that excites uniformly the whole granule. If a gradient of fluorescence has been imprinted, the bleach depth (fraction of fluorescence lost) measured by TIR is greater compared to the bleach depth measured by EPI, as TIR selectively probes the TIR-photobleached region. The second method (TIR-FRAP) involves monitoring fluorescence recovery over time following photobleaching with TIR excitation. As TIR selectively photobleaches fluorophores proximal to the glass interface, fluorescence recovery occurs as distal fluorophores diffuse in to the bleached area.

**Figure 1.**
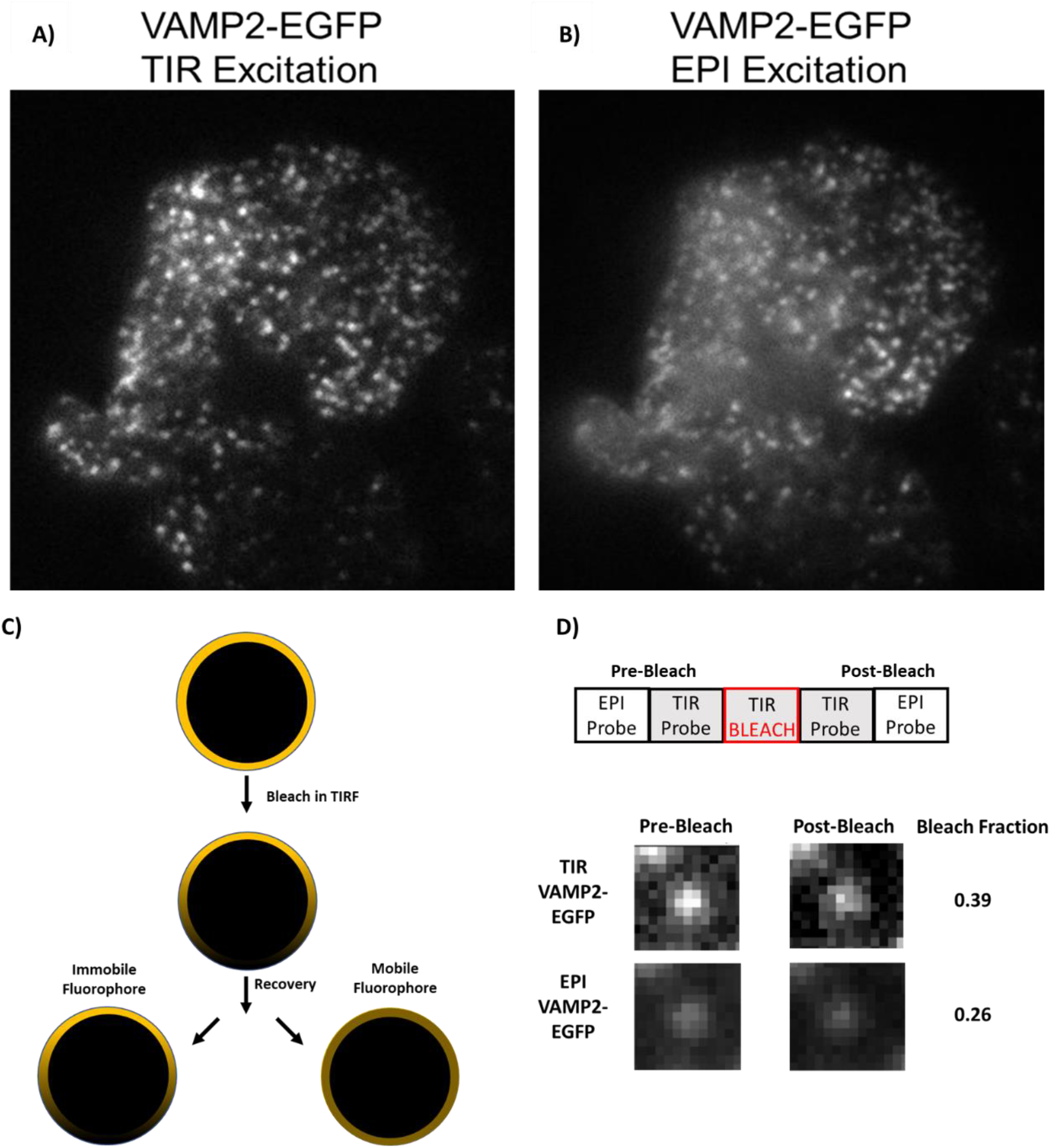
TIR based photobleaching protocol to measure the mobility of chromaffin granule membrane proteins. Image of a chromaffin cell expressing VAMP2-EGFP imaged using low-intensity TIR (**A**) or EPI (**B**) illumination. **C**) Schematic of the principle underlying the TIR-FRAP method. Chromaffin granules labelled with a fluorophore-tagged membrane protein are photobleached using TIR excitation light. The gradient of bleached fluorophore imprinted on the granule membrane dissipates over time if the protein is mobile, or remains stable if the protein is immobile. **D**) Fluorescence of the granules is measured using low intensity TIR and EPI illumination before and after bleaching in TIR. Shown is an example of a typical granule labelled with VAMP2-EGFP. The bleach depth (fraction of fluorescence lost) in TIR and EPI is calculated (1-F(post-bleach)/F(pre-bleach)).

First, as described in the theory section above, we derive by simulation the expected results for highly mobile and relatively immobile fluorophores, and consider the influence of scattered light, which contaminates the evanescent field, on the expected results.

### The influence of scattering on TIR/EPI bleach depth measurements

For highly mobile proteins, the bleach depth (i.e., the fraction of fluorescence lost) is identical when measured by TIR or EPI illumination (**Fig. 2 A**, solid 45° line), as the rapidly diffusing proteins uniformly distribute on the surface of the granule during the photobleaching step. For slowly diffusing proteins, the bleach depth as measured by TIR is relatively higher compared to the bleach depth measured by EPI illumination (**Fig. 2 A**, dotted line), as TIR selectively probes the photobleached region. However, in experimental settings, the evanescent field is not pure and is contaminated by scattered light originating either from the microscope optics (11, 12) and from refractive index irregularities in the cell (13). Previous measurements using a 1.65 numerical aperture lens and fluorescent beads estimated that ∼10% of the evanescent field is contaminated by scattered light at the coverslip/sample interface (14). Given the heterogenous refractive index of the cell imaged in the evanescent field (13), we expect that the degree of scattering to be greater than 10%. Thus, we derived the expected result for fluorophores on a surface of a hypothetical sphere with a D of 3 × 10^−10^ cm^2^/s photobleached in evanescent field containing 0, 10% or 20% scattered light at the coverslip/sample interface (**Fig. 2 A**, dotted and dashed lines). This value of D is within the range predicted by the recovery data (see below). As scattered light is not spatially selective, it photobleaches fluorophores uniformly on the surface of the granules, and decreases the difference between the bleach depths measured by low intensity TIR and EPI illumination and influences the sensitivity of our measurements. We then simulated the diffusion of fluorophores with varying D with 20% scattering present in the evanescent field (**Fig. 2 B**). Scattering reduced but did not eliminate the difference between TIR and Epi bleach depths.

**Figure 2.**
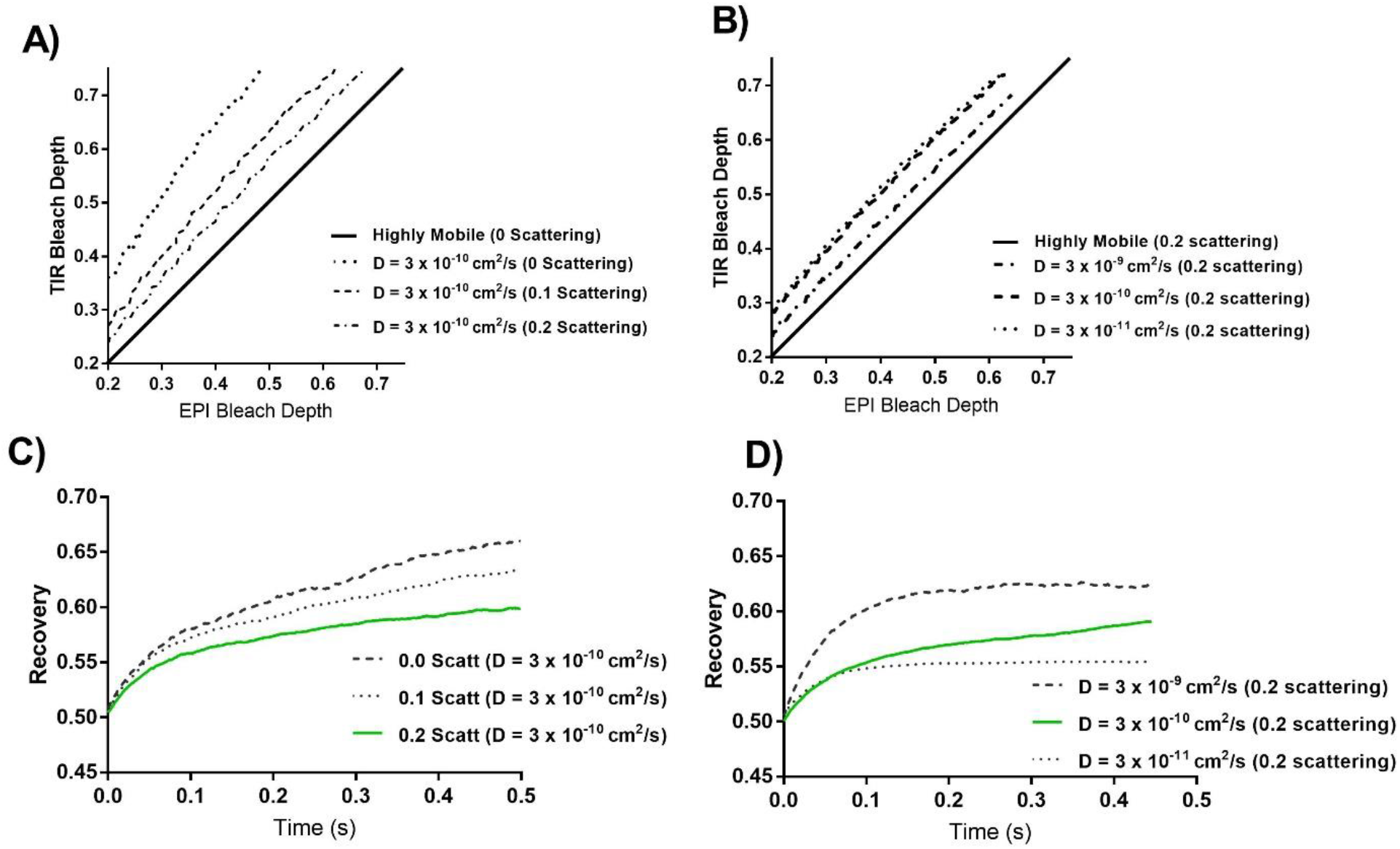
Simulation of diffusion on the surface of a sphere. Fluorescent proteins with a given diffusion coefficient (D), present on the surface of a 300 nm sphere, were photobleached for 45 ms in an evanescent field with a decay constant of 80 nm. The bleach depth (fraction of fluorescence lost) evident by TIR and EPI illumination (**A-B**), and fluorescence recovery following photobleaching (**C-D**) is plotted. The degree of scattering present in the evanescent field at the coverslip/sample interface was varied (**A, C**) while D was kept constant, or the degree of scattering was kept constant at 20 % while varying D (**B, D**).

### The influence of scattering on TIR-FRAP measurements

Similar to the TIR/EPI bleach depth simulations described above, we simulated time-dependent fluorescence recovery following TIR photobleaching and either varied the degree of scattering (0, 10%, or 20%) present in the evanescent field while considering a single D of 3 × 10^−10^ cm^2^/s (**Fig. 2 C**), or varied D while including 20 % scattering (**Fig. 2 D**). The presence of scattered light was found to decrease the initial rate of fluorescence recovery, and to a greater extent, reduced the extent to which fluorescence recovered (**Fig. 2 C**). Nonetheless, despite a substantial degree of scattering being present in the evanescent field, fluorescence recovery was strongly sensitive to D, with the initial rate of recovery and extent of recovery increasing with higher D (**Fig. 2D**).

Based on the simulations just described, both the TIR/EPI bleach depth measurements and TIR-FRAP recovery measurements are expected be sensitive to both scattering and diffusion. We included 20 % scattering (see above) while estimating diffusion coefficients based on our experimental data.

### TIR-fluorescence recovery after photobleaching (FRAP) measurements of fluorescent VAMP2, Syt1, and Syt-7 in chromaffin granule membranes in living cells

Primary bovine chromaffin cells were transfected with plasmids encoding secretory granule membrane proteins VAMP2-EGFP, Syt1-EGFP or msfGFP, or Syt7-EGFP, or the granule lumenal probe ss-mOxGFP (signal sequence of neuropeptide y fused to mOxGFP). Four to five days after transfection, cells were transferred to a physiological salt solution (PSS), and were photobleached using high intensity TIR excitation light to selectively photo bleach fluorophores proximal to the glass interface (**Fig. S 1**). The granule membrane and lumenal proteins were photobleached for 46 and 169 ms, respectively.

For each of the proteins examined, the bleach depth (fraction of fluorescence lost) as seen by TIR and EPI illumination following a TIR bleach is displayed in scatter plots (**Fig. 3 A – D**), and each data point represents an individual granule. For rapidly diffusing proteins (for which the bleached gradient is almost completely dissipated by the time of the probe measurement), the bleach depths as probed by either TIR or EPI illumination are expected to be equal to each other (indicated by the solid 45 ° black line); this was found to be the case for granules containing the granule lumenal probe ss-mOxGFP (signal sequence of neuropeptide y fused to mOxGFP) (**Fig. 3 D**). If proteins are immobile or slowly diffusing, then the bleach depth as probed by TIR is expected to be greater relative to the bleach depth probed by EPI (because the TIR probe selectively illuminates just the region that is bleached). Indeed, this was found to be the case for granules labelled with fluorescently tagged VAMP2, Syt1, and Syt7 (**Fig. 3 A – C**). This difference between the granule lumenal probe and membrane proteins is readily apparent when we compare the ratio of TIR/EPI bleach depths (**Fig 3. E**), which cluster close to 1 for ss-mOxGFP, indicating equivalent bleach depths, but are greater than 1 for VAMP2, Syt1, and Syt7 (i.e, greater bleach depth evident in TIR relative to EPI). We compared Syt1 with EGFP fused to its N terminus which is inside the granule lumen, or Syt1 with msfGFP fused to the C terminus which is in the cytoplasm. We did not observe a difference between the two fusion proteins, as the TIR probed bleach depth was greater relative to the EPI probed bleach depth in both cases, indicating that the result was not sensitive to the location of the fluorophore.

**Figure 3.**
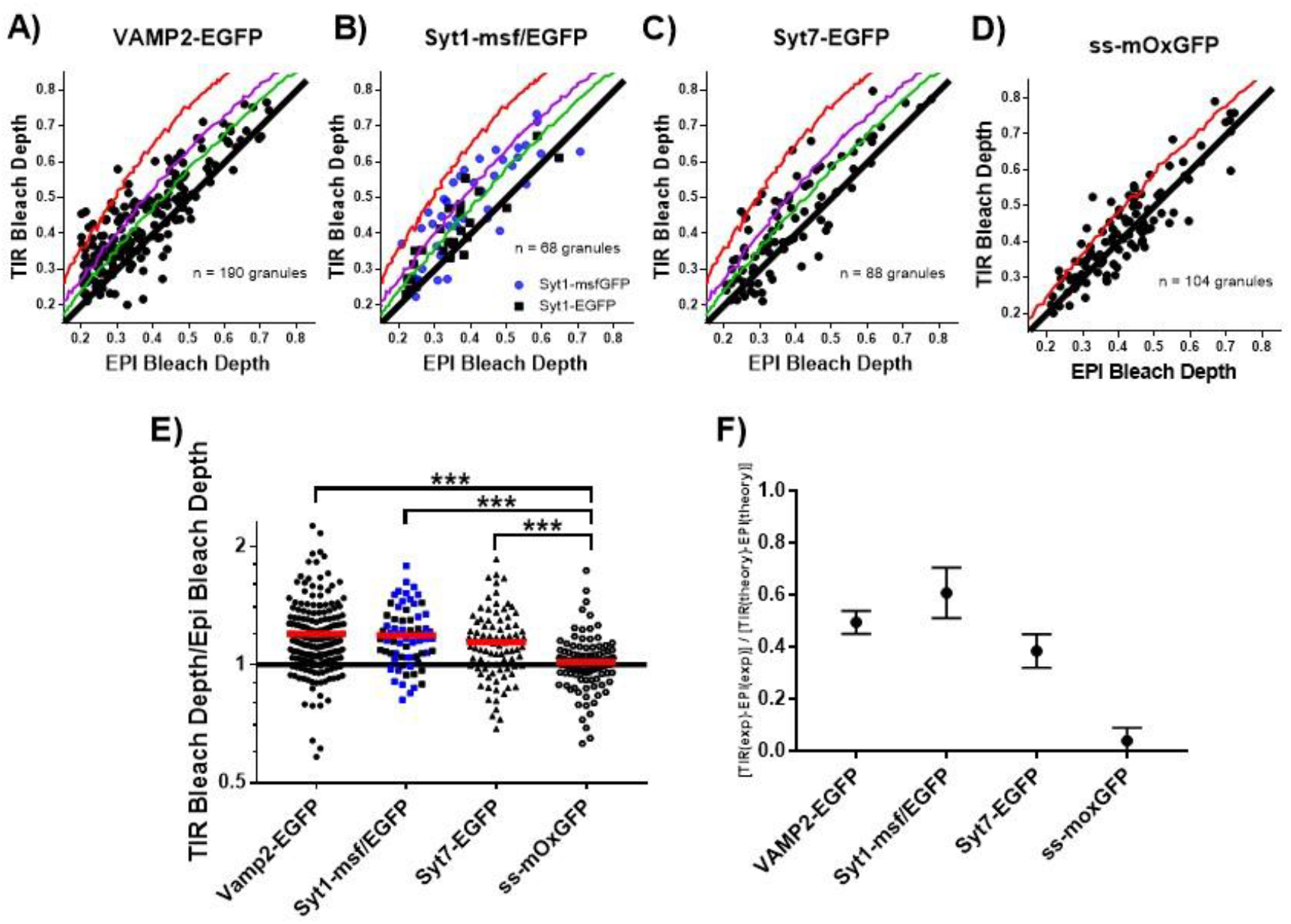
VAMP2 and synaptotagmins have low enough mobility in the granule membrane to retain a bleaching imprint within the duration of a bleaching pulse. Chromaffin cells were transfected with plasmids encoding **A)** VAMP2-EGFP, **B)** Syt1-msfGFP or Syt1-EGFP **C)** Syt7-EGFP, and **D)** the signal sequence of NPY fused to mOxGFP (ss-mOxGFP). Chromaffin cells expressing the fusion proteins were photobleached for 46 ms (granule membrane probes), or 169 ms (ss-moxGFP) with high intensity 488 nm excitation light in TIR mode. Fluorescence intensity was probed pre- and post-bleach using low intensity TIR and EPI illumination as described in the Methods section. The bleach depth (fraction of fluorescence lost) evident in TIR and EPI is plotted (1-F(pre-bleach)/F(post-bleach), and each data point represents an individual granule. The black 45° line is indicative of equivalent bleaching in EPI and TIR and is expected for highly mobile fluorophores. The red, purple and green curved lines are the expected result for a fluorophore with a diffusion coefficient of D = 3 × 10^−10^ cm^2^/s, photobleached in an evanescent field with 0 %, 10 % or 20 % scattering, respectively, and an exponential decay constant of 80 nm. These theoretical curves for membrane proteins (A-C) and for luminal proteins (D) are somewhat different, as discussed in the Theory section. **E)** The data shown in A-D expressed as the ratio of the bleach depths in TIR and EPI, the red line represents the median value in each group. Data from Syt1-msfGFP and Syt1-EGFP are colored blue and black, respectively. A ratio of 1 represents equivalent bleaching in TIR and EPI, *** indicates P < 0.0001 following a unpaired student’s t-test. **F)** The ratio of experimentally observed differences in bleach depths probed with TIR and EPI illumination compared to the theoretically expected differences for an immobile fluorophore. A ratio of 0 or 1 are expected for highly mobile or immobile fluorophores, respectively. Scattering was assumed to be 0 when deriving the expected theoretical results.

Utilizing the simulations described above, we compared the experimentally observed differences in bleach depths probed with TIR and EPI illumination to the theoretically expected differences for highly mobile or immobile fluorophores (**Fig 3. F**). A ratio of 0 or 1 are expected for highly mobile or immobile fluorophores, respectively. We found that the experimental data for ss-mOxGFP correlates closely with the expected result for highly mobile fluorophores, whereas the data from VAMP2-EGFP, Syt1-EGFP, and Syt7-EGFP granules has ratio between 0.4 and 0.6, indicating that they are relatively immobile. The theoretical results for this comparison were derived assuming 0 % scattering. Thus, VAMP2-EGFP, Syt1-EGFP, and Syt7-EGFP are less mobile than indicated by the ratio.

To verify that the greater fractional bleach depth evident by a TIR probe is due to photobleaching with TIR illumination, we repeated the same experiment but used high intensity EPI illumination to photobleach fluorophores. Bleaching in EPI does not imprint a gradient of photobleached fluorophore since EPI illumination uniformly photo bleaches fluorophores throughout the granule because it is not spatially selectively. As expected, the bleach depth as seen by TIR and EPI illumination following an EPI bleach is equivalent for all membrane and lumenal proteins examined (**Fig. S 1**).

Considered together, these experiments in living chromaffin cells indicate that TIR-based photobleaching selectively imprints a gradient of fluorescence, which is long lasting and readily apparent for granules labeled with the membrane proteins VAMP2, Syt1, and Syt7 fused to EGFP, but rapidly dissipates for the granule lumenal protein ss-mOxGFP. Thus, granule membrane proteins examined are relatively immobile compared to ss-mOxGFP. (It should be noted that another granule lumen protein, tissue plasminogen activator, is relatively immobile and sustains an imprinted gradient of fluorescence (15)).

### Time dependence of recovery

Following photobleaching with high intensity TIR excitation, we observed a time-dependent fluorescence recovery (**Fig. 4 A – C**). The data from individual granules were grouped based on the initial bleach depth immediately after the bleach (see Theory). The apparent recovery was ∼ 10% of the bleach fraction for granules expressing VAMP2-EGFP or Syt7-EGFP, and ∼ 5 % of the bleach fraction for granules expressing Syt1-EGFP or Syt1-msfGFP. As the surface of the granule is a closed system, and there is not an infinite pool of fluorophores available as in traditional TIR-FRAP experiments, recovery is expected to be much less than 100%, even in the case of high mobility.

**Figure 4.**
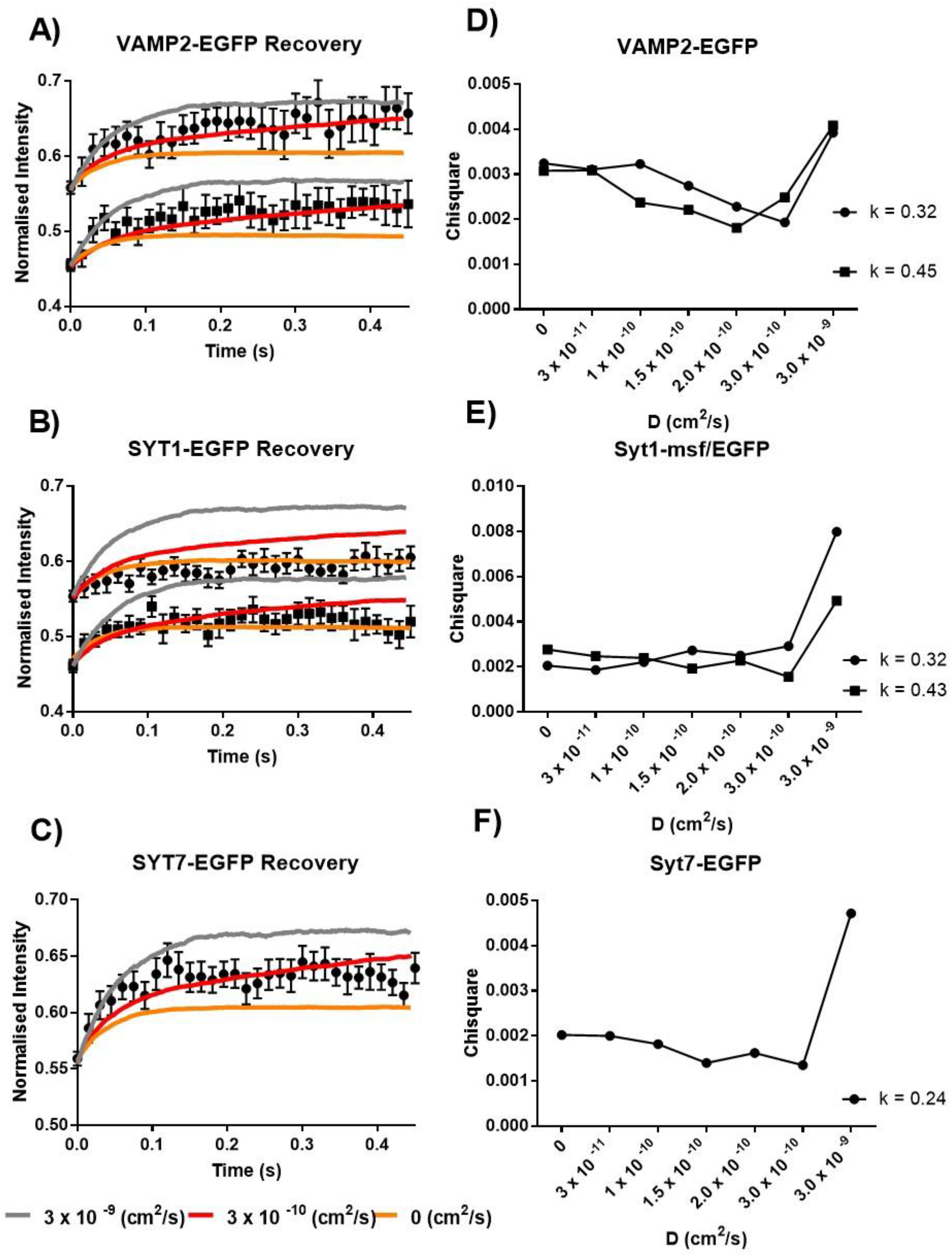
FRAP analysis of chromaffin granules expressing fluorophore tagged membrane or lumenal proteins. Chromaffin cells expressing **A)** VAMP2-EGFP, or **B)** Syt1-msf or EGFP, or **C)** Syt7-EGFP were photobleached with high intensity 488 nm light in TIR mode, and fluorescence recovery was measured using low intensity 488 nm light. Fluorescence recovery after photobleaching is shown. Experimental data were grouped into bins based on the fractional bleaching evident immediately after the photobleaching (bins: 0.4 – 0.5, 0.5 – 0.6, and each bin contains averaged data from 10 – 22 granules). Simulated theoretical curves are shown overlaid on the experimental data for three different diffusion coefficients. The kinetics and pathway (pathway 3) for irreversible bleaching were incorporated into the simulations. **D-F)** Chi-square goodness of fit analysis was used identify the simulated recovery curves that best described the experimental data.

It was anticipated that diffusion of the fluorophores in the granule membrane accounted for the experimentally observed recoveries. However, the recovery could also reflect, at least in part, the kinetics of reversible bleaching wherein the fluorophore is bleached but recovers instead of being irreversibly lost (16, 17). This issue is considered next.

### Reversible bleaching

In order to determine the extent of reversible bleaching, EPI rather than TIR illumination was used to bleach fluorescent granule membrane proteins. Fluorescence recovery was measured with low intensity EPI excitation (**Fig. 5**). As EPI illumination is not spatially selective, any observed fluorescence recovery reflects fluorophores that underwent reversible photobleaching. Reversible bleaching was, in fact, significant and comparable to the recovery from bleaching in the TIR-FRAP experiments. Reversible bleaching was approximately 5% of the pre-bleach fluorescence when bleaching reduced the fluorescence to 65% or 57% of the prebleach level (**Fig. 5**). The half-time for recovery was approximately 50 ms.

**Figure 5.**
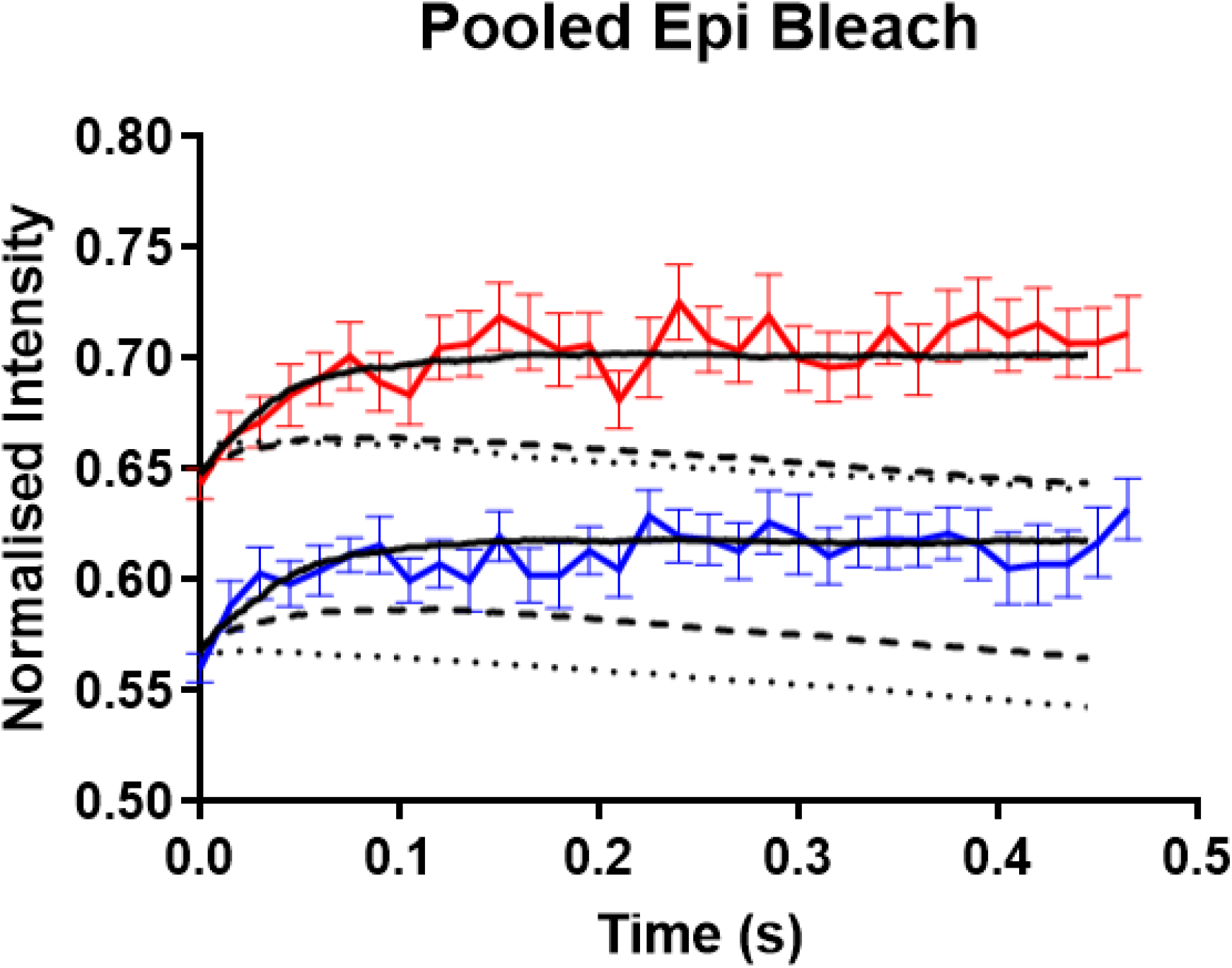
Fluorescence recovery of EGFP from a reversibly bleached state. **A)** Chromaffin cells transfected with plasmids encoding VAMP2-EGFP, Syt1-EGFP, or Syt7-EGFP were photobleached with high intensity 488 nm EPI excitation light. Fluorescence recovery following photobleaching was probed using low intensity EPI illumination. The data from cells expressing the various fusion proteins were pooled based on an initial bleach depth of either 50-60% (n = 19 granules), or 60-70% (n = 27 granules), and the normalized fluorescence recovery after the bleach is shown. The recovery reflects reversible bleaching. The final steady state fluorescence reflects irreversible bleaching. Three theoretically possible forms of reversible recovery were simulated: 1) reversible and irreversible bleaching occur in parallel (dark dashed line), 2) reversible bleaching protects against irreversible bleaching (light dashed line), and 3) irreversible bleaching can only occur from a reversibly bleached state (dark solid line). The third scenario best fit the experimental data.

To enable quantitative interpretation of the TIR bleaching experiments, we modeled three possible photochemical pathways for reversible and irreversible bleaching: 1) irreversible and reversible photobleaching occur in parallel. Any single fluorophore may undergo either one or the other of those processes; 2) a reversible bleaching event protects against irreversible bleaching; and 3) irreversible photo bleaching can only occur from a reversibly bleached state. In this third situation, a fluorophore has to encounter a minimum of two photons to be irreversibly photobleached: one to induce reversible photobleaching, and another to cause irreversible photobleaching. We found that only the third scenario adequately described the data (**Fig. 5**). In the first two modes of reversible recovery, there is initial fluorescence recovery followed by gradual irreversible photobleaching caused by low intensity probe illumination; however, in the third mode, the extent of recovery from a reversibly bleached state is high enough to compete with the rate of gradual irreversible photobleaching caused by probe illumination.

### Estimation of diffusion coefficients of granule membrane proteins

In order to determine diffusion coefficients, we generated a series of simulated TIR-FRAP recovery curves based upon diffusion on the surface of a sphere of the same diameter as a chromaffin granule (300nm). Diffusion coefficients were varied and the results compared to the experimental TIR-fluorescence recovery data. The kinetics and pathway (pathway 3) for irreversible bleaching were incorporated into the simulations. Scattering was assumed to account for 20% of the total bleaching intensity at the glass interface (see above). A diffusion coefficient of 0 represents recovery observed solely from a reversibly bleached state. We used chi-square-based goodness of fit analysis to determine the best fit (i.e., the diffusion coefficient that best described the data) in an unbiased way. The initial rising phase of recovery from VAMP2-EGFP and Syt7-EGFP expressing granules was best described by diffusion coefficients of 1.5 – 3 × 10^−10^ cm^2^/s (**Fig. 4. D, F**). Syt1-EGFP data could not be distinguished from a diffusion coefficient of 0 (**Fig. 4. E**), indicating that recovery observed from Syt1-EGFP expressing granules was likely only from a reversibly bleached state.

In summary, the observed rates of fluorescence recovery after photobleaching are strongly influenced by reversible bleaching, whose kinetics limits the ability to estimate low diffusion coefficients. The simulations indicate the diffusion constant for these granule membrane proteins is likely to be 3 × 10^−10^ cm^2^/s or less. A diffusion coefficient of 3 × 10^−9^ cm^2^/s or greater results in a suboptimal fit. The low mobilities indicated by fluorescence recovery are consistent with the ability of the fluorescent proteins to maintain the imprint of the fluorescent gradient imposed by bleaching in TIRF revealed by TIR/EPI bleach depth measurements (**Fig. 3**). The implications of these mobilities of the three proteins on the kinetics of exocytosis are considered in the Discussion.

## DISCUSSION

The granule membrane proteins VAMP2 (a v-SNARE) and Syt 1 and Syt 7 (Ca^2+^ sensors) are proposed to interact with lipids and cognate SNAREs on the plasma membrane to enable regulated exocytosis. As discussed below, the reactions that lead to fusion of an individual granule with the plasma membrane require multiple copies of these proteins. Thus, their diffusion to the fusion site could facilitate the fusion reaction and play role in fusion pore expansion. In this study we measured the mobility of these proteins in the membrane of chromaffin granules in living cells using a TIRF-based technique that is not limited to the optical resolution of the microscope. We found that two of the proteins (VAMP2-EGFP and Syt7-EGFP) have low (but non-zero) mobilities, and the third (Syt1-EGFP) has a mobility indistinguishable from zero. We consider the implications of this finding on regulated exocytosis in the following.

### Diffusion of VAMP2, Syt-1, and Syt-7 on chromaffin granule and plasma membranes

The *in situ* mobility of membrane proteins varies by almost 2 orders of magnitude depending upon the subcellular localization (18-21). Golgi proteins have diffusion constants of ∼5 × 10^−9^ cm^2^/s whereas plasma membrane proteins generally have the lowest mobilities with diffusion coefficients of approximately 10^−10^ cm^2^/s. Indeed, the plasma membrane SNARE protein syntaxin has been shown to exhibit mobilities ranging from 3.9 × 10^−10^ to 2 × 10^−11^ cm^2^/s (22). The low mobilities of plasma membrane proteins reflect interaction with intracellular cytoskeletal proteins and extracellular matrix proteins (23-27). Transfected VAMP2 and Syt1 disperse into the plasma membrane upon fusion, allowing the estimation of apparent diffusion constants (in the plasma membrane). VAMP-2-GFP diffuses from the site of exocytosis in the plasma membrane with an apparent diffusion constant of 2 × 10^−9^ cm^2^/s (28), similar to rhodopsin (29). Transfected Syt1-pHluorin diffuses into the plasma membrane with an apparent diffusion constant of 6 × 10^−11^ cm^2^ /s (30), similar to endogenous plasma membrane proteins. The slow diffusion of Syt1 may reflect the slow rate of dissociation of the protein from the fused granule membrane as well as interaction with intracellular and extracellular components. Syt7 barely disperses upon fusion, possibly because a stable, narrow fusion pore restricts dispersion into the plasma membrane (30).

Secretory granule and synaptic vesicle sizes are close to or below the diffraction limit for light microscopy. We adapted a TIR-FRAP based approach in our study which provided sufficient resolution perpendicular to the glass interface to study the diffusion of fluorophore-tagged VAMP2, Syt1, and Syt7 on chromaffin granules membranes in live cells. VAMP2 diffusion in the chromaffin granule membrane, 1.5 – 3 × 10^−10^ cm^2^/s, was an order of magnitude slower relative to its post-fusion diffusion in the granule membrane. Syt7 had a mobility comparable to VAMP2 on the chromaffin granule membrane. Syt1 diffusion was too slow to be resolved, suggesting slower diffusion compared to VAMP2 and Syt1.

### VAMP2 and Syt diffusion and granule fusion-ability, rate of fusion, and fusion pore expansion

The assembly of multimeric complexes at the site of fusion depends upon the local availability of the component proteins. The minimum number of fusogenic proteins required at the granule-plasma membrane contact site to mediate the fusion reaction is unknown. Reconstitution-based biochemical analyses have suggested that as little as three SNARE complexes are sufficient to form a fusion pore (31-33). In hippocampal synapses, two copies of VAMP2 may be sufficient for synaptic vesicle fusion and it was suggested that diffusion of a VAMP2 molecule was necessary to consummate the fusion reaction (34). A study in isolated chromaffin cells found that at least three SNARE complexes are required for efficient exocytosis (35). There is disagreement whether synaptotagmin interaction with SNARE proteins is required to enable Ca^2+^-triggered fusion (36-39). There is no generally accepted model for Syt-SNARE interactions. Nonetheless, there is strong evidence of functional cooperation between Syt1 and VAMP2, and reconstitution-based studies have shown that Syt1 penetrates membranes while bound to SNARE complexes (40-42). It is plausible that an equal number of Syt1 and SNARE proteins need be present at the fusion site.

We calculated the minimum time required for an equal number of VAMP2 and Syt 1 to reach an area encompassed by an early fusion pore of 2 nm diameter, given an initial density of VAMP2 and Syt 1 reported to be present on synaptic vesicles (43, 44). We estimate that ∼ 1 copy of VAMP2 and ∼ 0.3 copies of Syt1 will already be present on average in such an area. In order to estimate the time required for diffusion of 4 copies of VAMP2 and Syt 1 to the nascent fusion pore, we modelled the granule membrane as an infinite plane containing VAMP2 and Syt 1 at densities reported on synaptic vesicles, and modelled the fusion pore as a “perfect sink” that instantly and permanently traps molecules that it encounters ((Eq. 5.79 in Crank (45) **Fig 6 A**). Because this method does not take into account the depletion of the finite pool of molecules in the granule membrane, it estimates the *minimum* time required. We reasoned that 4 copies would be sufficient to enable both fusion pore opening and subsequent expansion. We found that it would take ∼ 7 and 55 ms for 4 copies of VAMP2 and Syt 1, respectively, to arrival at the nascent fusion pore (**Fig 6 A**).

**Figure 6.**
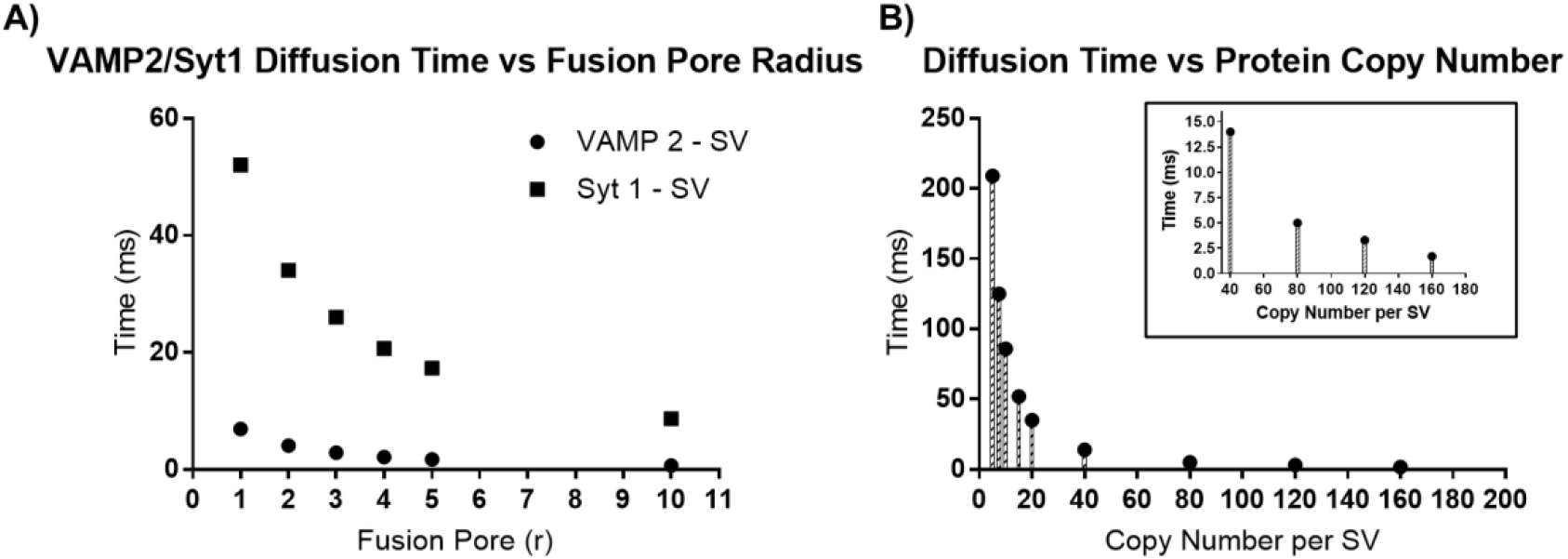
Influence of diffusion and protein abundance on the time required for proteins to reach a fusion pore of a defined radius. **A)** Time required for 4 copies of Syt1 or VAMP2 to arrive at a fusion pore of a defined radius. A diffusion coefficient of 3 × 10^−10^ cm^2^/s was used to describe the mobility of both VAMP2 and Syt1, and the concentration of VAMP2 and Syt1 was set to 70 and 15 copies per synaptic vesicle, respectively. The diffusion time calculations were performed as described in the discussion section. **B)** The influence of vesicle membrane protein abundance on the time required for proteins to reach a fusion pore. Given a D of 3 × 10^−10^ cm^2^/s, we calculated the amount of time needed for 4 copies of a protein to reach a fusion pore 1 nm in radius, with protein abundance ranging from 5 – 160 copies/synaptic vesicle which corresponds to a density of 820 – 26000 copies/µm^2^. Inset shows a magnified view of the 40 – 160 copy number/vesicle range.

Because the copy numbers of VAMP2 and Syt 1 on large dense core secretory granules are unknown, we investigated the effects of different protein densities on the time required for 4 copies of a protein with a D of 3 × 10^−10^ cm^2^/s to reach a 1 nm radius fusion pore (**Fig 6 B**). As expected, diffusion times increased as protein abundance deceased. For example, if there were 7 copies of Syt 1 on synaptic vesicles instead of 15, the time required for 4 copies to reach a nascent fusion pore would be increased from ∼ 55 ms to ∼ 125 ms. Stochastic variation in protein abundance on individual vesicles is likely to result in variable diffusion times to a nascent fusion pore, a variable that could impact the range of latencies before fusion is consummated.

What is the likelihood that the assembly of the fusion complexes would be limited by the rate of diffusion of the granule proteins to the fusion site? This depends upon the pathway for the initiation of the fusion reaction. In one model, granules are thought be in a stable interaction with the plasma membrane through partially zippered trans-SNARE interactions with Ca^2+^ triggering the fusion reaction. In this scenario, VAMP and synaptotagmin would already be assembled at the fusion site and the fusion reaction would not be diffusion-limited. This trans-SNARE complex has been difficult to detect and its existence has been questioned (2). In a second model, the interaction of the v- and t-SNARES is all or none and very rapid, without trans SNARE pre-engagement. It is driven by a large downhill energy gradient for SNARE complex formation (2). There is evidence for this possibility. FRET measurements suggest that the plasma membrane t-SNARE acceptor complex of syntaxin and SNAP25 for granule VAMP2 forms 100 ms before fusion (46-48). Additionally, granules continue to jitter as much as 100 nm (much greater than the ∼ 10 nm length of the tetrahelical SNARE complex) within 100 ms of fusion (4, 5). Both these observations argue against the formation of trans-SNARE complexes 100 ms before fusion. If the latter model is correct, then formation of the initial fusion pore could be limited by the diffusion times of VAMP2 and/or synaptotagmin (approximately 7 ms and 50 ms, respectively) to the fusion site.

It is interesting to consider the time delay to membrane fusion following a stimulus. Following elevation of intracellular Ca^2+^ in chromaffin cells, there is delay of ∼10 – 100 ms prior to the detection of membrane fusion events by capacitance or amperometric measurements, following by a rapid burst of secretion (49-52). The initial rapid phase of exocytosis has both fast and slow components which time constants ranging from ∼1-100 ms and ∼10-1000 ms (49), respectively. These rates vary further depending on the source and type of preparation, with chromaffin cells in adrenal slices displaying faster release kinetics relative to dissociated cells in culture (53). These delays to fusion following a stimulus roughly correspond to the diffusion time required for multiple copies of VAMP2 and Syt1 to reach the fusion site. Thus, in addition to the time required for Ca^2+^ to reach fusion-ready granules, the diffusion of fusogenic proteins to the vesicle-plasma membrane contact site could contribute to the observed delays to fusion following a stimulus.

Could protein diffusion on synaptic vesicles affect the kinetics of synaptic transmission? It would seem unlikely because of latencies as short as 60μs for exocytosis following the influx of Ca^2+^ (54). It is often assumed that the short latency reflects an intrinsically rapid fusion reaction. On the other hand, the short latency could be a statistical manifestation of a large number of release-ready synaptic vesicles in tens to hundreds of boutons, each containing ∼ 10 docked vesicles per active zone that on average have a much slower fusion rate (55). In this scenario, the 60 µs latency reflects a stochastic selection of dynamic and rare synaptic vesicles with VAMP2 and synaptotagmins perfectly aligned with t-SNARES on the plasma membrane (56). Milliseconds later in the response, vesicles with more characteristic slower fusion rates would be manifest.

Protein diffusion may play another role in exocytosis. Both fusion pore stability and the subsequent rate of fusion pore expansion in reconstituted fusion reactions increase with increasing numbers of SNARE complexes (32, 57). Thus, diffusion of VAMP2 and synaptotagmin into the nascent fusion pore may influence both its stability and expansion.

In summary, while the molecular machinery required to mediate regulated exocytosis has been identified, the details of protein and lipid interactions that enable membrane fusion and fusion pore expansion are not well understood. This study demonstrates that VAMP2, Syt 1 and Syt 7 in the granule membrane have relatively low mobilities that may influence the dynamics of these processes.

## MATERIALS AND METHODS

### Chromaffin cell culture and transfection

Bovine chromaffin cells were isolated and transfected using the Neon Transfection System (Thermo Fisher Scientific) as previously described (58, 59). For transfection of GFP-tagged granule membrane proteins, 1 ug of the plasmid encoding the fusion protein was used per 10^6^ cells. All plasmids used were verified by DNA sequencing, and details regarding the promoter, and linker sequences are provided in the supplement (**Table S. 1**). Imaging experiments were performed in physiological salt solution (PSS) containing 145 mM NaCl, 5.6 mM KCl, 2.2 mM CaCl_2_, 0.5 mM MgCl_2_, 5.6 mM glucose, and 15 HEPES, pH 7.4. All live-cell imaging experiments were performed at 34°C 4–6 d after cell isolation.

### TIR-FRAP microscopy and image analysis

Prismless (through-the-objective) TIR excitation was obtained by directing a laser beam from a 100 mW solid-state (488 nm) laser (Coherent OBIS, CA, USA) onto a computer-controlled galvanometer mirror and then toward a side pore of an Olympus inverted microscope. The aligned excitation beam was focused and positioned near the periphery of the back focal plane of a 60× 1.49-NA, oil immersion objective (Olympus) so that the laser beam was incident on the coverslip at ∼70° from the normal giving a decay constant for the evanescent field of ∼80 nm. The galvanometer mirrors were computer controlled though a DAQ board (National Instruments) and a custom LabView program, and allowed for rapid switching between TIR and EPI excitation by modulation of the incidence angle. To allow for rapid switching between high-intensity bleach and low-intensity probe excitation, the OBIS laser was operated in analog modulation mode and the input voltage was controlled by the same custom LabView program that controlled the galvanometer mirrors. Fluorescence emission was collected through the 1.49-NA objective, and images were acquired using a CMOS camera (Prime, Photometrics).

Two different but related types of TIR/EPI data were gathered: “bleach depth” and “FRAP". For TIR/EPI bleach depth measurements, single sets of EPI and TIR images were utilized, one set immediately before and one set immediately after the bleach pulse which lasted 46 ms (as shown in **Fig 1.B**). For TIR and EPI-FRAP measurements, a time sequence of numerous TIR or EPI images were utilized both before and after the bleach pulse. For both types of measurements, images were acquired at a rate of 65 hz. The images were subsequently processed using ImageJ (Fiji distribution) and data analysis was performed using custom programs written in Python and Interactive Data Language (IDL). Granules harboring GFP-tagged proteins were identified as bright puncta and changes in fluorescence intensities were monitored over time within a defined circular region of interest (ROI) that encompassed the whole puncta. Background intensity was measured from a circular region immediately adjacent to the granule, and subtracted from the granule fluorescence intensity. Only granules with ROI average intensity at least 15 % greater than background intensity throughout the movie (i.e, before and after photobleaching) were included in the final analysis. In the TIR and EPI bleach depth experiments, the emission intensities from individual granules were averaged over a set of three successive frames to reduce shot noise. For the TIR and EPI-FRAP experiments, emission intensities were used without any averaging.

## Acknowledgements

We thank Drs. Kevin P. Bohannon and Mary A. Bittner for many helpful discussions. msfGFP used to construct the C-terminal tagged syt1-msfGFP fusion protein was a gift from Dr. Benjamin Glick (University of Chicago). This work was supported by NIH Grant R01-170553 to RWH and DA. E.R.C. is an Investigator of the Howard Hughes Medical Institute, and acknowledges support from NIH grants MH061876 and NS097362.

## FIGURES

**Supplemental Figure 1.**
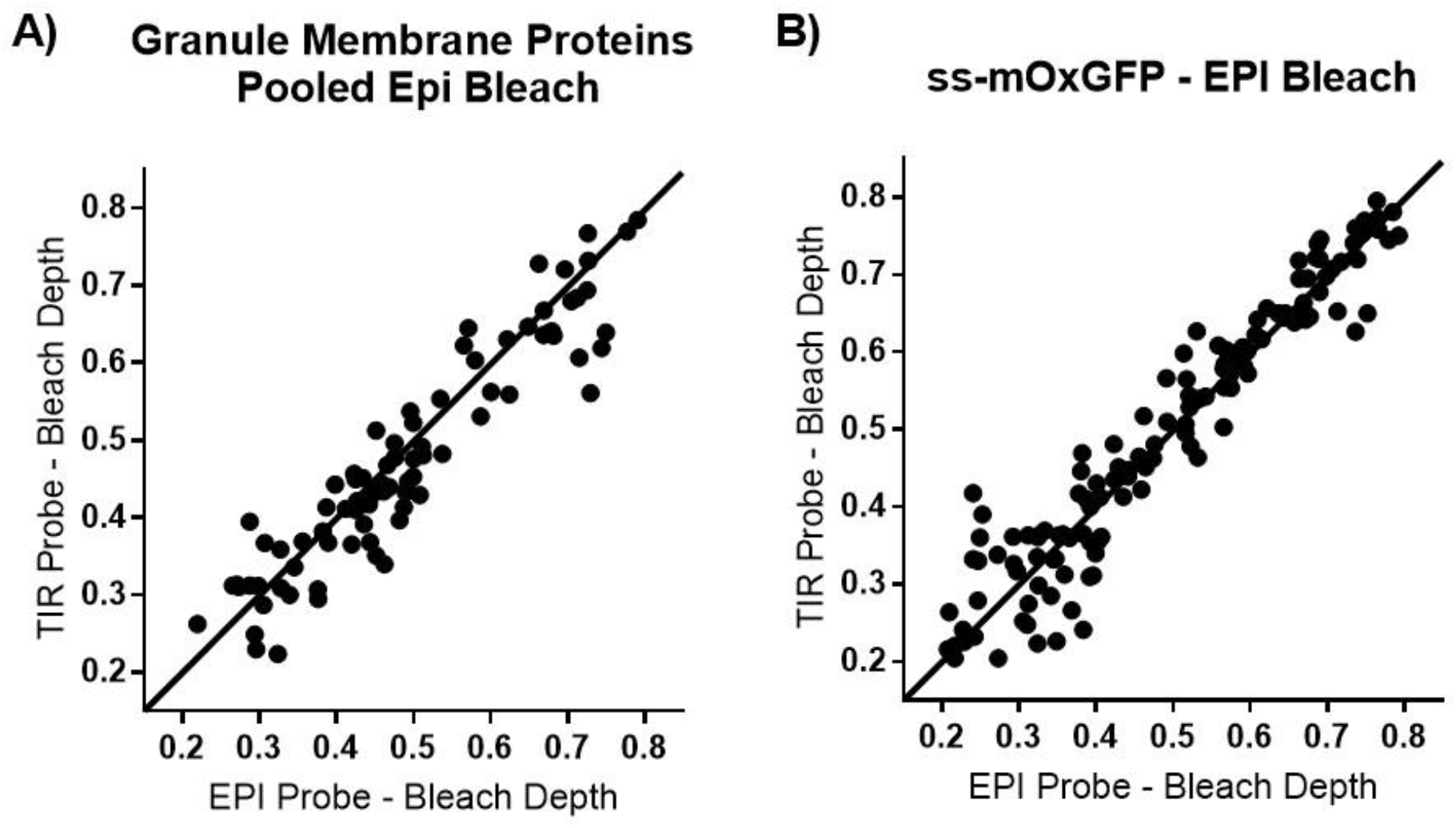
Photobleaching with EPI does not imprint a gradient of fluorescence on the granule. **A)** Chromaffin cells were transfected with plasmids encoding VAMP2-EGFP, Syt1-msfGFP or Syt1-EGFP, Syt7-EGFP or the lumenal probe **B)** ss-mOxGFP and photobleached with high intensity 488 nm excitation light in EPI. Fluorescence intensity was probed pre- and post-bleach using low intensity TIR and EPI illumination. The bleach depth (fraction of fluorescence lost) evident in TIR and EPI is plotted (1-F(pre-bleach)/F(post-bleach). Each data point represents an individual granule. The black 45° indicates equivalent bleaching in EPI and TIR.

**Supplementary Table 1.**
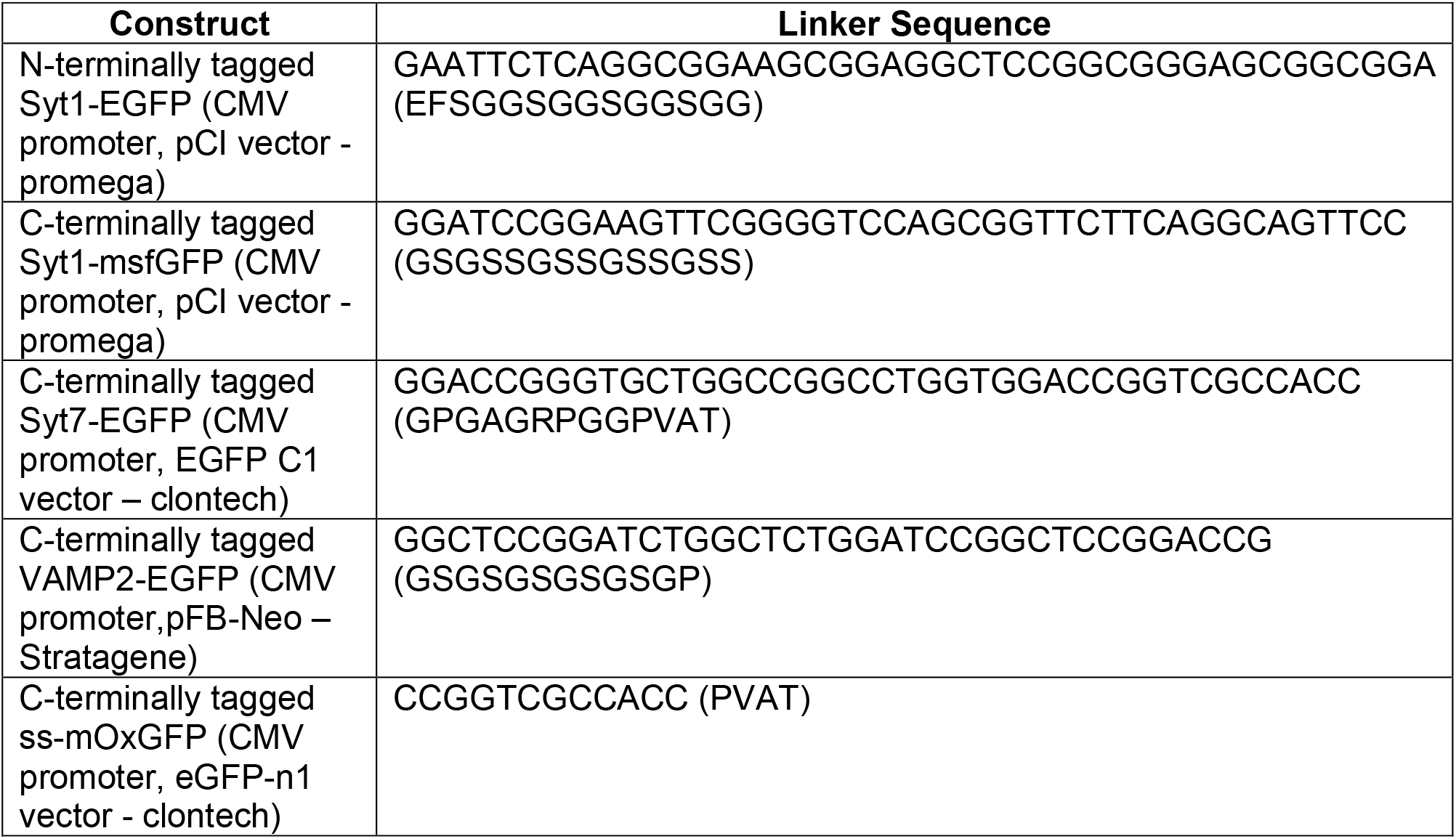
Promoter, vector, and linker sequences of all of the protein expression constructs used in the study.

